# Endocrine aryl hydrocarbon receptor signaling is induced by moderate cutaneous exposure to ultraviolet light

**DOI:** 10.1101/487355

**Authors:** Babak Memari, Loan Nguyen-Yamamoto, Reyhaneh Salehi-Tabar, Michela Zago, Jorg Fritz, Carolyn J. Baglole, David Goltzman, John H. White

## Abstract

Links between solar UV exposure and immunity date back to the ancient Greeks with the development of heliotherapy. Skin contains several UV-sensitive chromophores and exposure to sunlight can produce molecules, such as vitamin D3, that act in an endocrine manner. We investigated whether the aryl hydrocarbon receptor (AHR), an environmental sensor and ligand-regulated transcription factor activated by numerous planar compounds of dietary or environmental origin, could be the target of endocrine photoproducts produced by cutaneous UV exposure. 15-to 30-minute exposure of cells to a minimal erythemal dose of UVB irradiation *in vitro* induced translocation of the AHR to the nucleus, rapidly inducing site-specific DNA binding and target gene regulation. Importantly, *ex vivo* studies with *Ahr* wild-type or null fibroblasts showed that serum from mice whose skin was exposed to a 15 min UVB dose, but not control serum, contained agonist activity within 30 min of UV irradiation, inducing AHR-dependent gene expression. Moreover, a 15-min cutaneous UVB exposure induced AHR site-specific DNA binding and target gene regulation *in vivo* within 3-6 hr post-irradiation in blood and in peripheral tissues, including intestine. These results show that cutaneous exposure of mice to a single minimal erythemic dose of UVB induces rapid AHR signaling in multiple peripheral organs, providing compelling evidence that moderate sun exposure can exert endocrine control of immunity through the AHR.

## Introduction

Links between solar ultraviolet (UV) irradiation and immunity date back millennia; the concept of heliotherapy was developed by the ancient Greeks, and reintroduced in the 19^th^ century with the advent of the sanatorium movement in Europe to treat consumption (tuberculosis) (1). UV light, which is divided into UVC (100–290 nm), UVB (290–320 nm), and UVA (320–400 nm) (2), is filtered by the ozone layer such that all UVC and much of the UVB radiation does not reach the earth’s surface. This, depending on latitude and the angle of the sun, leaves a variable portion of UVB and most UVA unfiltered. Skin contains numerous UV-sensitive chromophores, leading to the production of several photoproducts upon exposure that can regulate both the innate and adaptive arms of the immune system (2). Much attention in this regard has focused on understanding the role of vitamin D_3_, produced cutaneously from UVB-induced photo- and thermal conversion of 7-dehydrocholesterol, as an immune system regulator (3,4). However, there is evidence that exposure of skin to UVB at levels that induce minimal or no increase in circulating vitamin D metabolites leads to immune system regulation (5). While UV exposure can modulate immunity by altering antigen presentation or cytokine levels (6), other endocrine factors may also be produced by photochemical reactions in exposed skin.

One attractive candidate for such endocrine signaling is the aryl hydrocarbon receptor (AHR), a ligand-regulated transcription factor highly expressed in barrier organs of the body (7,8). While it was initially characterized for its capacity to bind dioxin (2,3,7,8-tetrachlorodibenzodioxin), a potent and metabolically-resistant environmental toxicant, the AHR functions physiologically as a dietary or environmental sensor that can recognize a wide array of largely planar molecules of environmental origin (8–10). The AHR is a member of the basic helix-loop-helix-PAS (bHLH-PAS) transcription factor family. It heterodimerizes with its homologue ARNT (AHR nuclear translocator) to bind cognate DNA sequences called xenobiotic, dioxin, or AHR response elements (XREs, DREs, or AHREs, respectively). The AHR can also heterodimerize with RELB, an anti-inflammatory and immune-modulatory subunit of the transcription factor NF-κB (7). One of the most strongly induced AHR target genes encodes CYP1A1, which hydroxylates and metabolically inactivates several physiological AHR ligands (11).

Importantly, the AHR has emerged as a key regulator of innate immunity (8–10), and its signaling can be induced locally in skin by UVB exposure (12). In this study, we tested the hypothesis that exposure to a single minimal erythemic dose of ultraviolet B radiation can induce systemic AHR signaling *in vivo*. We show that UVB exposure *in vitro* induces nuclear localization of the AHR within 15 min, leading to its site-specific DNA binding and target gene regulation. A series of *ex vivo* and *in vivo* experiments with mice revealed that cutaneous exposure to a single dose of UVB leads to release of AHR agonists into the circulation, and to induced AHR DNA binding and target gene regulation *in vivo* in peripheral tissues within 3hr after exposure. Taken together, these studies show that UV exposure of skin rapidly induces endocrine signaling through the AHR.

## Results and Discussion

To test for effects of single doses of moderate UV exposure on AHR signaling, we used an irradiation protocol that generated 1.2 kJ/m^2^ or 2.5 kJ/m^2^ after 15 or 30 min exposures, which is equivalent to approximately 1-2 minimal erythemal doses (13). We used a narrow-band (311nm) UV source for *in vitro* studies because broadband UVB induces elevated levels of cell death *in vitro* even after limited exposure (14,15). Other work has shown that the level of irradiation used produces no or only moderate increases in circulating levels of vitamin D metabolites *in vivo* (5). We carried out control experiments *in vitro* in order to test the kinetics of UV-induced nuclear translocation of AHR, using a protocol that generated complete separation of nuclear lamin and cytoplasmic actin (Fig. S1A). Under these conditions, a single 10-30 min exposure induced AHR nuclear translocation (Figs. 1A, B) to a degree that was similar to that induced by AHR agonist FICZ (Fig. 1C). In other control experiments, a single dose of UV had no effect on the subcellular localization of the vitamin D receptor after 1 hr, although addition of the AHR ligand precursor kynurenine led to accumulation of the AHR in the nucleus over the same period (Figs. S1B, C), indicating that there was not a general effect on nucleocytoplasmic shuttling arising from UV exposure. To probe further the effects of UVB exposure on AHR activation, we analyzed expression of the well-characterized AHR target gene, *CYP1A1*, 4 hr after 15 min UVB exposure, which revealed an induction that was approximately 2-fold higher than that induced by FICZ after the same period (Fig. 1D). UVB exposure also induced *CYP1A1* expression in THP-1 macrophages, 4 hr after exposure (Fig. S3A), and single dose led to sustained *CYP1A1* expression in SCC25 epithelial cells (Fig. S3B). The effects of treatment on AHR target gene regulation were substantiated further by Western blot analysis of CYP1A1 protein, which revealed an increase in expression over a 24 hr period after 15 min of UVB exposure (Fig. 1E). The role of the AHR in UVB-stimulated gene expression was confirmed by depletion of the receptor in SCC25 cells, which abolished induced expression of *CYP1A1* and AHR target gene *IL1B*. These studies were performed using either a single siRNA targeting *AHR* (Fig. 1F) or pooled siRNAs recognizing completely independent sites on the transcript (Fig. S3C), with similar results. A similar knockdown experiment in epithelial cells abolished induction of *CCL1* and *S100A9* (Fig. S3D), previously identified as target genes (16). Finally, we used chromatin precipitation (ChIP) assays to test the effects of UV exposure on AHR DNA binding to previously characterized XREs in the promoters of the *CYP1A1*, *IL10*, and *IL23A* genes, all of which, along with AHR target gene *IL22*, are inducible by AHR agonists or by UVB (Figs. 1D, Figs. S3E,F) (16). UVB exposure for 15 min enhanced AHR DNA binding to XREs of target promoters in both epithelial and myeloid cells (Figs. 2A, B).

**Figure 1:**
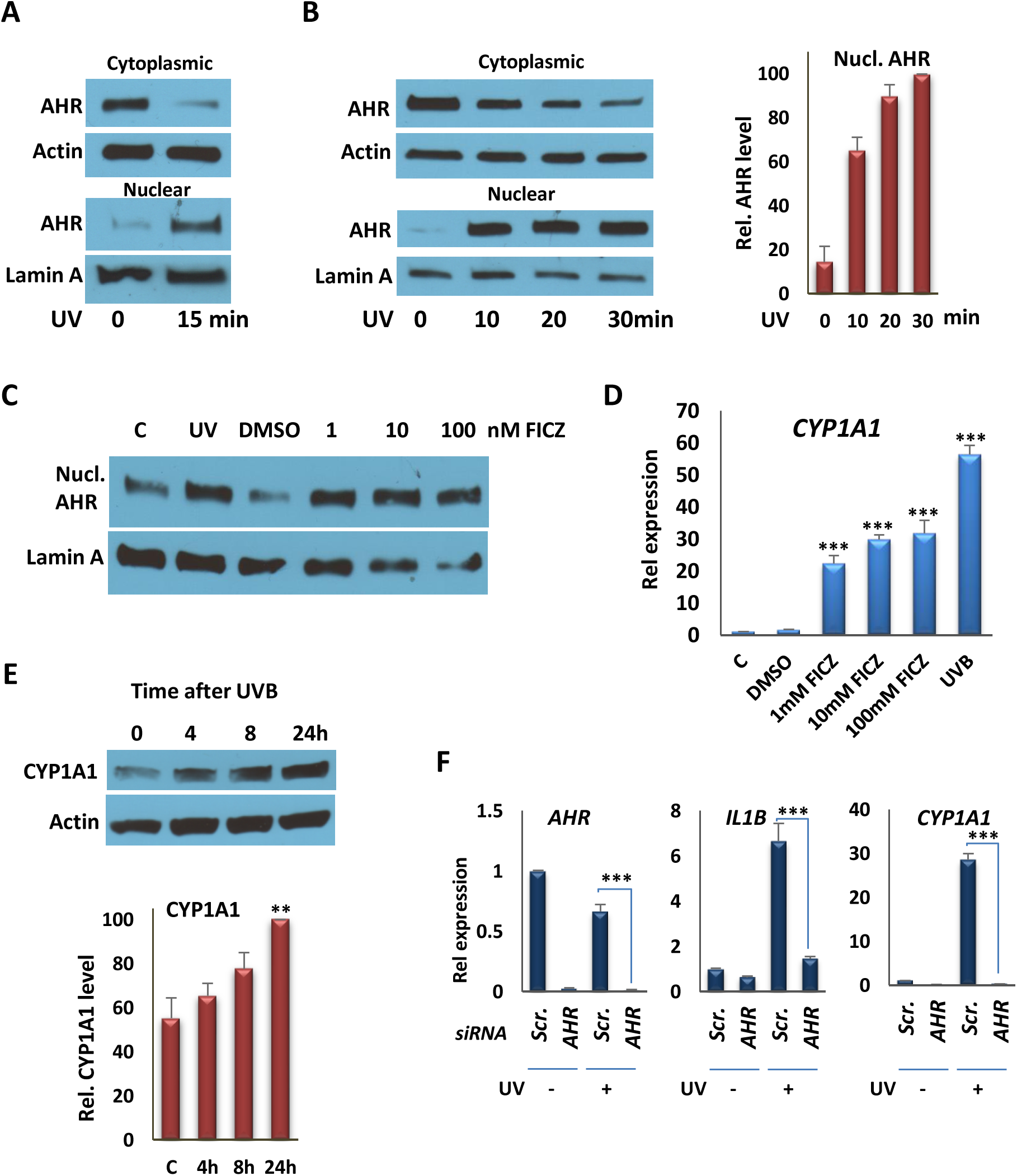
UVB induces AHR translocation to the nucleus and the expression of its target genes, *in vitro*. **A)** Western Blot analyses of AHR from nuclear and cytoplasmic fractions from SCC25 cells 1 hr following irradiation with 15 min UVB. Actin was probed as a cytoplasmic marker; lamin A was probed as a nuclear marker. AHR and the internal controls were taken from the same blot. Blot images are provided in the Supplementary Fig S4A. **B)** (left panel) Western Blot analyses of AHR in nuclear and cytoplasmic fractions from THP-1 cells 1 hr following irradiation with UVB (10, 20, or 30 min). (right panel) The quantification of Western Blot analyses, represented in the left panel. AHR and the internal controls were taken from the same blot. Blot images are provided in the Supplementary Fig S4B. **C)** Western Blot analyses of nuclear AHR 1 hr following irradiation with UVB (15 min) or FICZ (1, 10, or 100 nM) treatment. AHR and the internal controls were taken from the same blot. Blot images are provided in the Supplementary Fig S4C. **D)** RT-qPCR analysis of *CYP1A1* transcription in SCC25 cells 4 hr after exposing cells to UVB for 15 min or treating them with FICZ, as indicated. **E)** Effect of a single 15 min UVB exposure on production of CYP1A1 protein in SCC25 cells 4, 8 or 24 hr after exposure. (Upper) Western blot of a single experiment. (Lower) Quantification of results of three independent experiments. CYP1A1 and Actin were taken from the same blot. Blot images are provided in the Supplementary Fig S4D. **F)** RT-qPCR analysis of *AHR*, *CYP1A1* and *IL1B* expression in SCC25 cells 4 hr after exposing cells to UVB for 15 min following knockdown of the *AHR* gene with siRNA #1. **P ≤ 0.01, ***P ≤ 0.001 as determined by one-way ANOVAs followed by Tukey’s post hoc test for multiple comparisons.

**Figure 2:**
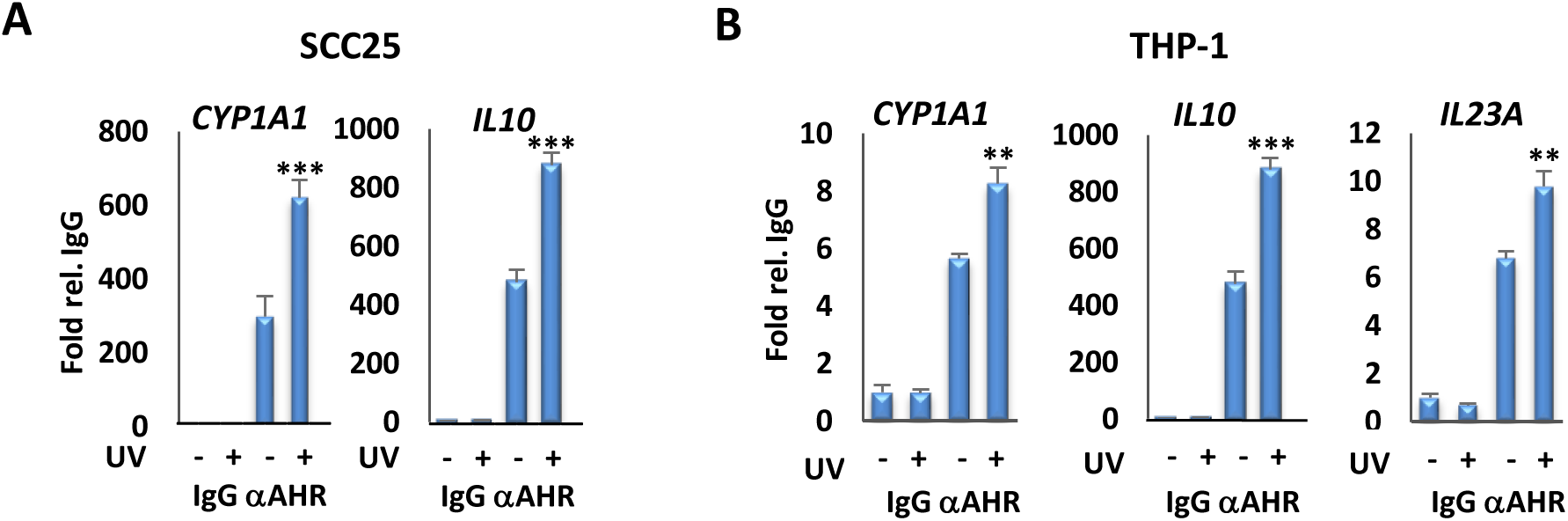
UVB induces AHR recruitment to the promoter of target genes. **A)** Analysis of AHR recruitment to the XRE motifs of the *CYP1A1* and *IL10* promoters by ChIP assays, followed by qPCR, in SCC25 cells 4 hr after irradiation with UVB for 15 min. **B)** Analysis of AHR recruitment to the XRE motifs of promoters of the *CYP1A1*, *IL10* and *IL23A* genes by ChIP assays followed by qPCR, in THP-1 cells 4 hr after irradiation with UVB for 15 min. **P ≤ 0.01, ***P ≤ 0.001 as determined by one-way ANOVAs followed by Tukey’s post hoc test for multiple comparisons.

Collectively, the *in vitro* experiments above show that moderate UVB exposure induces AHR signaling within minutes. Previous work has shown that cutaneous UV exposure in mice can induce AHR function in skin (12). To test for the effects of a single, moderate UVB dose on AHR signaling in mice, we used a broadband UV source (peak at 313nm), which is more reflective of the composition of sunlight than narrowband UV. Initial controls showed that a single exposure of mice to UVB for 15 min (1.2 kJ/m^2^) induced expression of AHR target genes *Cyp1a1*, *Il22* and *Il23a* in skin (Fig. S2), as expected (12). To extend these findings, and test whether AHR ligands are released into the circulation after exposure of mouse skin to UVB, serum was extracted from male or female control or UVB-exposed animals 30 min after 15 min of irradiation, and incubated with primary fibroblasts derived from wild-type or *Ahr*-null animals. Incubation with serum from irradiated but not control mice induced AHR target gene expression in wild-type but not *Ahr*-null fibroblasts (Fig. 3A). Furthermore, the effect in wild-type fibroblasts was completely blocked by the AHR antagonist CH223191 (Fig. 3A). These experiments show that moderate UVB exposure of skin leads to rapid (within 30 min) release into the circulation of agonists that induce robust AHR target gene expression, consistent with the kinetics of AHR induction seen above *in vitro*. This effect is dependent on the presence of the receptor but independent of the sex of mice providing either donor serum or recipient fibroblasts. In addition, incubation of sera from UVB-exposed but not control animals with naïve T cells induced expression of Th17 markers *Il17a* and *Il17b* (Fig. 3B), consistent with the capacity of AHR agonists to induce Th17 cell differentiation (17).

**Figure 3:**
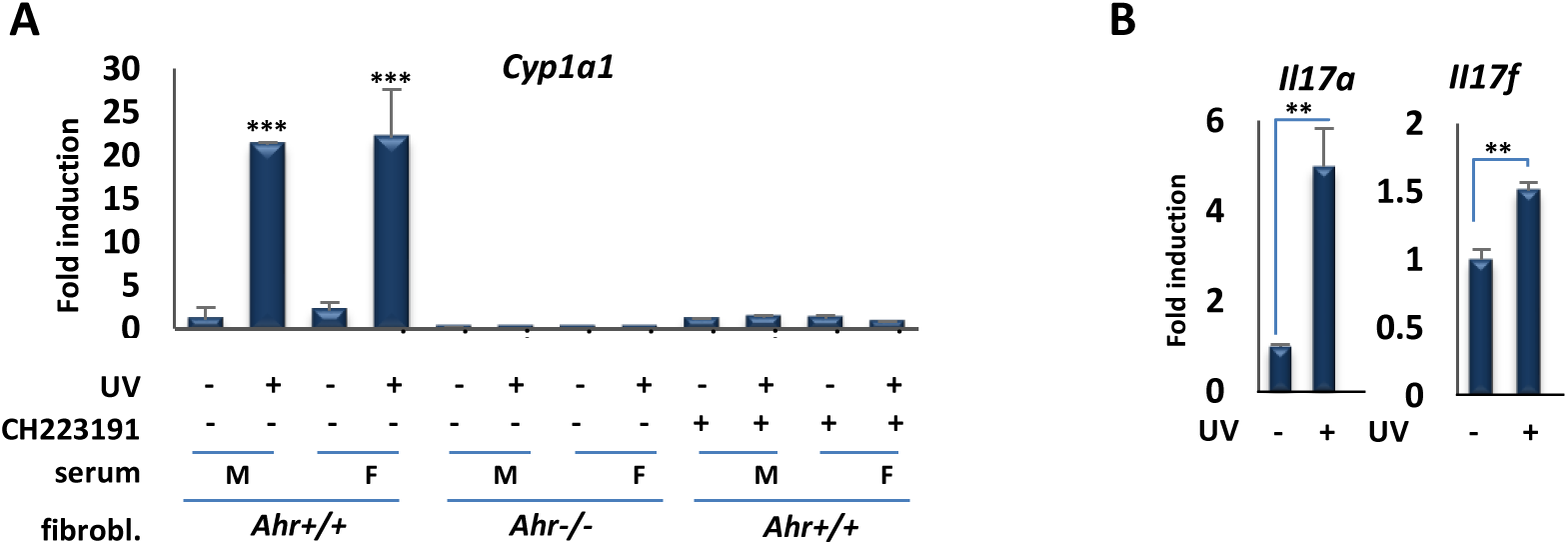
Serum of mice exposed to moderate UVB induces the expression of AHR target genes, *ex vivo*. **A)** Serum of UVB-exposed mice induces the expression of AHR target genes in mouse fibroblast. RT-qPCR assay of *Cyp1a1* mRNA expression in primary fibroblasts derived from wild-type or *Ahr*-null mice 6 hr following treatment with the serum of control unexposed (-) mice or UV-exposed (+) male or female mice and AHR inhibitor, CH223191. Serum obtained from mice 30 min after irradiation with (1.2 kJ/m2) UVB for 15 min. **B)** *Il17a* and *Il17b* gene expression following incubation of naïve T cells with sera from UVB-exposed mice. **P ≤ 0.01, ***P ≤ 0.001 as determined by one-way ANOVAs followed by Tukey’s post hoc test for multiple comparisons.

Taken together, studies above reveal serum derived from animals exposed to moderate levels of UVB induces robust AHR signaling *ex vivo*. To test for the effects of UV exposure on induction of AHR signaling in peripheral tissues *in vivo*, we determined the effects of a single 1.2 kJ/m^2^ dose of UVB on the expression of several receptor target genes in blood, liver or intestine, 3 or 6 hr after UVB exposure, as indicated (Figs. 4A-D). In all cases, induction of AHR target gene expression was observed, consistent with cutaneous UVB exposure inducing rapid, endocrine signaling. Target genes included *Il22* and *Il23a* in intestine, consistent with the effects of UVB on their expression *in vitro* (Fig. S3E,F). IL23 is a macrophage cytokine that is a target of AHR signaling in human and mouse (16), and part of an AHR-regulated cascade; its signaling leads to production of the innate immune cytokine IL22 in AHR-regulated innate lymphoid cells in the intestine (18). These results were strongly supported by ChIP analysis of AHR DNA binding to well-characterized XREs *in vivo*. In control experiments, a single UVB dose induced AHR binding to XREs in the *Cyp1a1* and *Il23a* promoters in skin (Fig. 5A). The same exposure also stimulated AHR DNA binding in small intestines 6 hr after UVB exposure (Fig. 5B).

**Figure 4:**
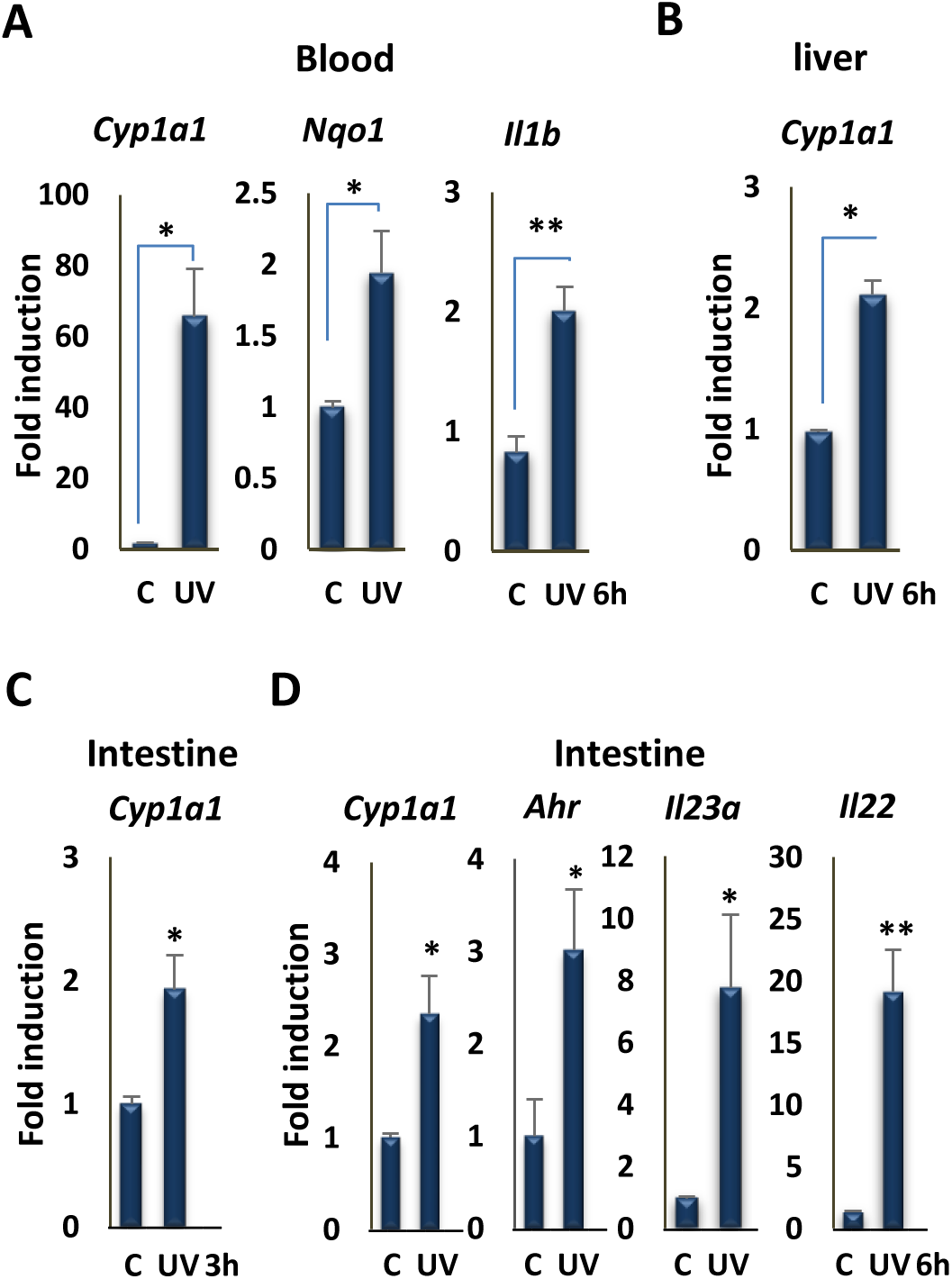
UVB irradiation induces the expression of AHR target genes in different tissues. **A)** RT-qPCR assay of *Cyp1a1*, *Nqo1*, and *Il1b* mRNA expression in blood of mice 6 hr following irradiation with 1.2 kJ/m2 UVB for 15 min (n = 3 per group). **B)** RT-qPCR assay of the *Cyp1a1* mRNA expression in liver of mice 6 hr following irradiation with UVB for 15 min (n = 3 per group). **C)** RT-qPCR assay of the *Cyp1a1* mRNA expression in intestine of mice 3 hr following irradiation with UV for 15 min (n = 3 per group). **D)** RT-qPCR assay of the *Cyp1a1*, *Ahr*, *Il23a* and *Il22* mRNA expression in intestine of mice 6 hr following irradiation with UVB for 15 min (n = 3 per group). *P ≤ 0.05, **P ≤ 0.01, as determined by one-way ANOVAs followed by Tukey’s post hoc test for multiple comparisons.

**Figure 5:**
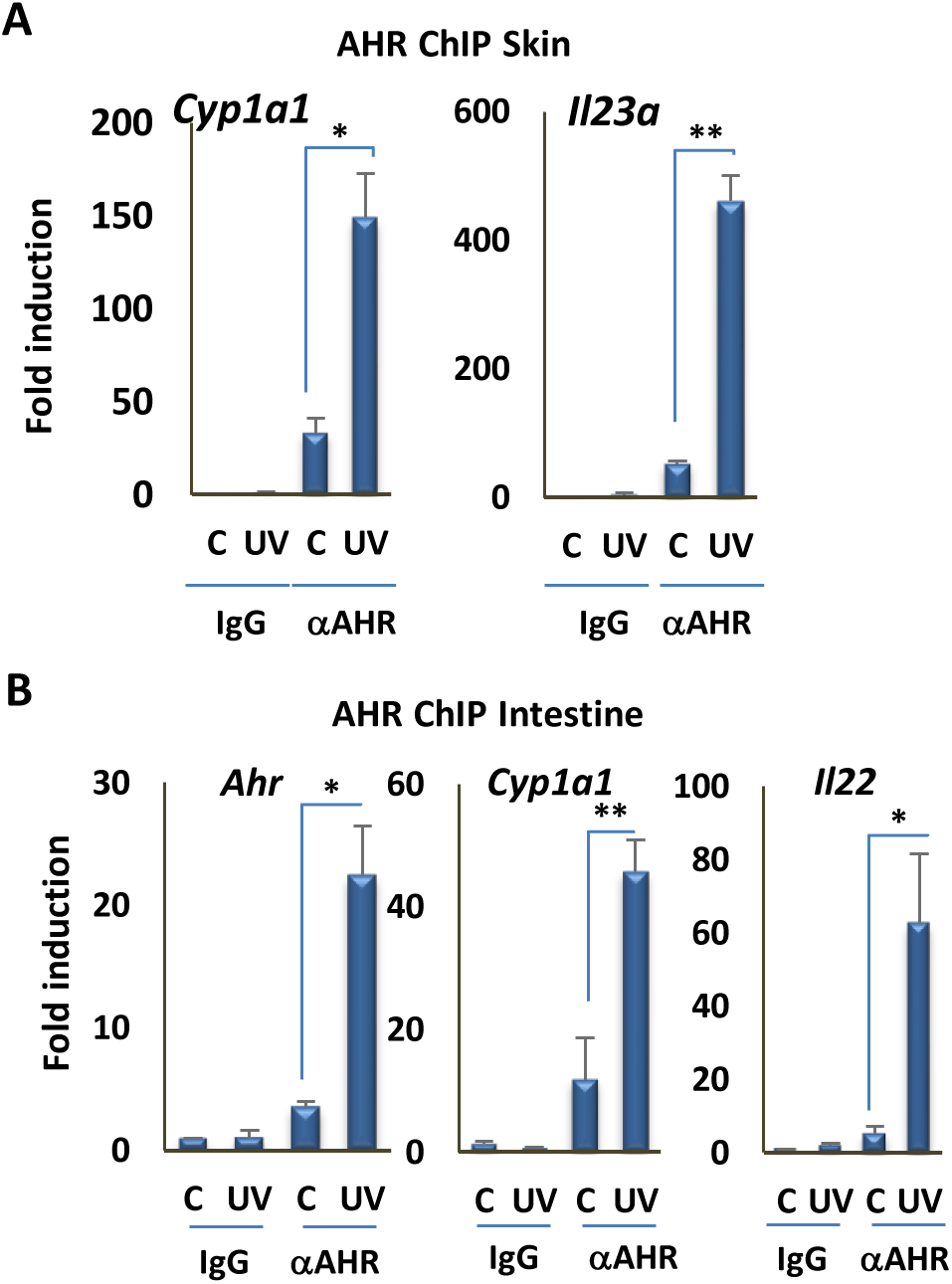
UVB irradiation induces the recruitment of AHR to the promoter of its target genes in different tissues. **A)** Analysis of AHR recruitment to XRE motifs of the *Cyp1a1* and *Il23a* promoters by ChIP assay, followed by qPCR, in skin of control and UVB-exposed mice (n = 3 per group). **B)** Analysis of AHR recruitment to the XRE motifs of the *Cyp1a1*, *Ahr*, and *Il22* promoters by ChIP assays followed by qPCR, in intestine of mice 6 hr following irradiation with UVB for 15 min (n = 3 per group). *P ≤ 0.05, **P ≤ 0.01, ***P ≤ 0.001 as determined by one-way ANOVAs followed by Tukey’s post hoc test for multiple comparisons.

The data presented above provide compelling evidence that a single exposure to as little as 15 min of moderate intensity UVB irradiation leads to rapid activation, nuclear translocation, DNA binding, and target gene regulation by the AHR *in vitro* and *in vivo*. Our findings are consistent with UV exposure generating sufficient levels of AHR agonist *in vitro* to induce nuclear translocation within 15 min. Similarly, *ex vivo* studies with serum from UV-exposed mice harvested 30 min after single-dose exposure showed the AHR agonist activity is released rapidly into the circulation following UV treatment. Notably, several indole-containing compounds can serve as AHR ligands (7,10). For example, the indole-containing amino acid tryptophan can produce AHR ligands via several routes, including photodimerization to produce FICZ, and condensation with cysteine to produce ITE [2-(1′H-indole-3′-carbonyl)-thiazole-4-carboxylic acid methyl ester] (19,20). In experiments conducted with peripheral tissues of UV-exposed mice, we observed induced AHR DNA binding and target gene activation within 3-6 hr in tissues including small intestine, an important site of AHR signaling (9,21). Regulatory events in small intestine included AHR binding to and induction of the direct target gene encoding IL22, an innate immune cytokine and a key component of AHR-regulated intestinal immunity, as well as induction of the gene encoding IL23A, whose signaling lies upstream of IL22. Taken together, these data provide compelling evidence that moderate cutaneous UVB exposure induces endocrine signaling through the AHR, a ligand-regulated environmental sensor. Given the emerging role of AHR signaling in the immune system in mice as well as humans, in particular in barrier organs, our data provide further mechanistic evidence for a role of cutaneous UVB exposure in the endocrine regulation of immunity.

## Materials and Methods

### Cells

SCC-25 cells (CRL-1628; ATCC) were cultured in DMEM/F12 (319-085-CL; Wisent) supplemented with 10 % FBS (non-heat inactivated). THP-1 monocytic cells (TIB-202; ATCC) were cultured as suspension cells in RPMI 1640 (350-005-CL, Wisent) with 10 % FBS (non-heat inactivated; Wisent) and differentiated into adherent THP-1 macrophages with phorbol myristate acetate (PMA; 100 ng/ml) for 24 hr. The *Ahr* knock-out mouse fibroblasts were cultured in EMEM (30-2003; ATCC) supplemented with 15 % FBS.

### Primary lung fibroblast cell culture

For isolation of primary lung fibroblast cells from C57BL/6 wild-type mice, the protocol described in (22) was used. Briefly, after sacrificing the mice, lungs were immediately transferred to the ice-cold PBS. Lung tissue was cut into ~1 mm pieces, washed with ice-cold PBS and placed in DMEM/F12 medium containing 0.14 wunsch units/mL “Liberase TM” and Pen-Strep was added and kept with gentle shaking at 37 °C in cell incubator for 30 min. Cells and tissue fragments were washed with DMEM/F12 + 15 % FBS, three times to inactivate liberase and cultured in DMEM/F12 (319-085-CL; Wisent) + 15 % FBS + Pen-Strep for up to 14 days. After attaching of the cells to the plate, medium was changed to EMEM (30-2003; ATCC) supplemented with 15 % FBS.

### Western Blot assay and antibodies

Cells were lysed by adding 300 μl of lysis buffer (0.5 % IGEPAL-CA-630 Sigma, 0.5 mM EDTA, 20 mM Tris-Base pH 7.6, 100 mM NaCl, protease inhibitor cocktail) into 10 cm plates. All plates were scraped and transferred into 1.5 ml vial on ice and sonicated in an ice bath for two cycles 10 sec, amplitude 30 % on the Vibra-Cell Sonics system VCX-750 (Four-element probe, 3 mm stepped microtip). All vials were centrifuged at 10,000 RPM for 10 min at 4 °C. The protein concentration was determined using Bio-Rad DC Protein assay. The normalized and denatured protein samples (containing Laemmli buffer, heated at 70 °C 10 min) were loaded on Precast Bio-Rad Mini-TGX 4–15 % polyacrylamide gel (456–1084). The following antibodies were used for immunoblotting: AHR (BML-SA210-0100; ENZO Lifesciences), Actin (sc-1615; Santa Cruz), CYP1A1 (sc-393979; Santa Cruz), and VDR (sc-13133; Santa Cruz).

### Nuclear and Cytoplasmic protein extraction

Cells were lysed in lysis buffer (0.5% Nonidet P-40, 0.5 mM EDTA, 20 mM Tris [pH 7.6], 100 mM NaCl, Sigma protease inhibitor cocktail), followed by centrifugation at 2000 RPM. The supernatant was collected as the cytoplasmic protein fraction. The cytoplasmic fraction in this protocol contains all cell components except nucleus. The pellet (nuclei) was dissolved in RIPA buffer (150 mM NaCl, 1.0% IGEPAL, 0.5% sodium deoxycholate, 0.1% SDS, and 50 mM Tris, pH 8.0) and sonicated at 35% power for 3 cycles of 10 sec, then centrifuged at 10000 RPM. The supernatant used as the nuclear fraction.

### Small interfering RNA–mediated knockdown

The following siRNAs were used for knockdown: negative control siRNA (non-silencing; QIAGEN; SI03650325); siRNA #1: Hs_AHR_6 Flexi Tube siRNA (QIAGEN; SI03043971). ON-TARGET plus non-targeting siRNAs (Dharmacon; D-001810-0X); siRNA #2: SMART pool ON-TARGET plus human AHR siRNA (Dharmacon; L-004990-00-0005). siRNA knockdowns were performed as described previously (16).

### RNA extraction

For cell culture, RNA extraction was initially performed by using 1ml TRIzol reagent or Tri-Reagent (Favorgen). Cells were homogenized by pipetting up and down several times and transferred into 1.5 ml vial. 200 μl chloroform was added, and the vials were shaken horizontally for 15 sec and kept for 10 min at room temperature and then centrifuged at 10,000 RPM for 10 min at 4 °C. Next, 400 μl of supernatant was passed through the first column (white) of a Blood/Cultured Cell Total RNA Mini Kit (FABRK Favorgen), and mixed with the same volume of 70 % ethanol, and passed through the RNA extraction second column (red). The RNA was washed and eluted according to the manufacturer’s Favorgen instructions. Tissues were frozen using dry ice or the flash frozen method immediately after sacrificing the mice. RNA was isolated from tissues by pulverizing with a Scienceware liquid nitrogen cooled mini mortar and pestle set, and about the size of a match head dissolved by pipetting in 1 ml TRIzol reagent or Tri-reagent (Favorgen). 200 μl chloroform was added, shaken horizontally for 15 sec, and kept at room temperature (RT) for 10 min. The remaining steps are similar to those for RNA extraction from cells. For blood, 200 μl of blood were mixed with in 1ml TRIzol-LS. Small intestine was used for RNA extraction from intestinal tissue.

### Quantitative PCR

cDNA was prepared from 500 ng of total RNA. Reverse transcription (RT) was performed with a Bio-Rad iScript cDNA Synthesis Kit (cat. no. 170-8891), and qPCR was performed using SsoFast EvaGreen Supermix (cat. no. 172-5211) on an Illumina Eco qPCR cycler. Primers used for analysis of gene expression are in table S1.

### Chromatin Immunoprecipitation (ChIP) assay

ChIP assays were performed as previously described (16). AHR Ab (sc-5579; Santa Cruz) and normal rabbit IgG (2729; Cell Signaling) were used for ChIP assays. Primer sequences used in qPCR for ChIP analysis are listed in Table S1.

### Mice

3-4 months-old C57BL/6 mice were used. All animal experiments were reviewed and approved by the McGill University Animal Care Committee.

### UV irradiation

The backs of the mice were shaved and 24 hr later mice were injected with ketamine hydrochloride and irradiated with broadband FS20T12/UVB Bulb (two lamps) from 40 cm distance for 15 or 30 min. For *in vitro* experiments, SCC-25 or THP-1 cells were cultured in 6 or 10 cm plates, and cells were irradiated in plates without lid under a biological hood using UVB narrowband TL20W/01-RS (two lamps) from 40 cm distance for 15 min or the times indicated in the figures. For cell culture experiments, only narrowband UVB lamps were used, because broadband lamps cause significant cell apoptosis and detachment. TL20W/01RS lamps emit a narrow peak around 311 nm, whereas FS20T12/UVB lamps emit a continuous spectrum from 275 to 390 nm, with a peak emission at 313 nm. Approximately, 65 % of that radiation is within the UVB wavelength range.

### Mouse Serum Preparation

Following UV irradiation, mice were sacrificed and approximately 1 ml blood was obtained by cardiac puncture. The blood was kept vertically in vial for 30 min at room temperature in the dark to allow the blood to clot in an upright position and no movement, and then centrifuged for 10 min at 3000 RPM. The serum was transferred to new vial. Centrifugation was repeated, for 5 min, in case of presence of blood or transparent viscous material in the supernatant (serum).

### Naïve T cell isolation from spleen

Naïve T cells were isolated by EasySepMouse Naïve CD4+ T Cell Isolation Kit # 19765 from mouse spleen. Mice were anesthetized by ketamine and spleens were transferred in cold Hanks’ Balanced Salt Solution containing 2% fetal bovine serum (FBS) and disrupted. All aggregates and debris were removed by passing the suspension through a 70 μm nylon strainer and centrifuged at 300 x g for 10 min and re-suspended at 1 x 10^8^ nucleated cells/mL in recommended medium. Antibody-based cell depletion was performed as per manufacturer’s recommendations.

### Th17 Polarization of Mouse CD4 Cells

Naïve T cells need activation for proliferation. For activation of naïve T cells, the protocol from BioLegend Inc. was used, as previously published (17). Bacterial plastic petri dishes (higher affinity for coating) were coated with anti-mouse CD3ε (clone 145-2C11, 2 μg/ml) and incubated at 37°C for 2 hr or 4°C overnight. All dishes were washed 3 times with sterile PBS. Then CD4 cells were added at 1 x 10^6^ /ml to the plates and cultured for 3 days in the presence of anti-mouse CD28 (clone 37.51, 5 μg/mL), IL-6 (50 ng/mL), TGF-β1 (1ng/mL), anti-mouse IL-4 (10 μg/mL), anti-mouse IFN-γ (10 μg/mL). Then non- and UVB-exposed-mouse serum were added into two separate plates. At day 3, cells were washed once and re-stimulated in complete medium with 500 ng/ml PdBU + 500 ng/mL ionomycin, in the presence of Brefeldin A for 4 hr. List of antibodies and cytokines is as follows: Anti-mouse CD3ε (clone 145-2C11, LEAF format, cat. # 100314), Anti-mouse CD28 (clone 37.51, LEAF format, cat. # 102112), Anti-mouse IL-4 (LEAF format, cat # 504108), Anti-mouse IFN-γ (LEAF format, cat. # 505812), Recombinant mouse IL-6 (carrier-free, cat. # 575702), Recombinant human TGF-β1 (carrier-free, cat. # 580702), Recombinant mouse IL-23 (carrier-free, cat. # 589002), Brefeldin A (cat. # 420601), PdBu (Phorbol 12, 13-dibutyrate, cat. # P1269 from Sigma) were used for Th17 Polarization.

### Statistical Analysis

All experiments are representative of three to five biological replicates. Statistical analysis was conducted using SYSTAT13 Trial by performing one-way ANOVA, followed by the Tukey test for multiple comparisons.

## Supporting information

## Acknowledgments

This work was supported by an operating grant from the Canadian Institutes of Health Research (CIHR, PJT-74294) to J.H.W., C.J.B., and J.H.F.

## Author Contributions statement

B.M conceived the project and performed the experiments. L.N.-Y. and B.M did the animal work. B.M., M.Z. and R.S.-T prepared the cells for experiments. J.H.W., J.F., C.J.B., D.G., and B.M. designed research and revised the manuscript. J.H.W., B.M., and R.S.-T analyzed data. B.M. and J.H.W. wrote the paper.

