## Supplementary material for "Endocrine aryl hydrocarbon receptor signaling is induced by moderate cutaneous exposure to ultraviolet light"

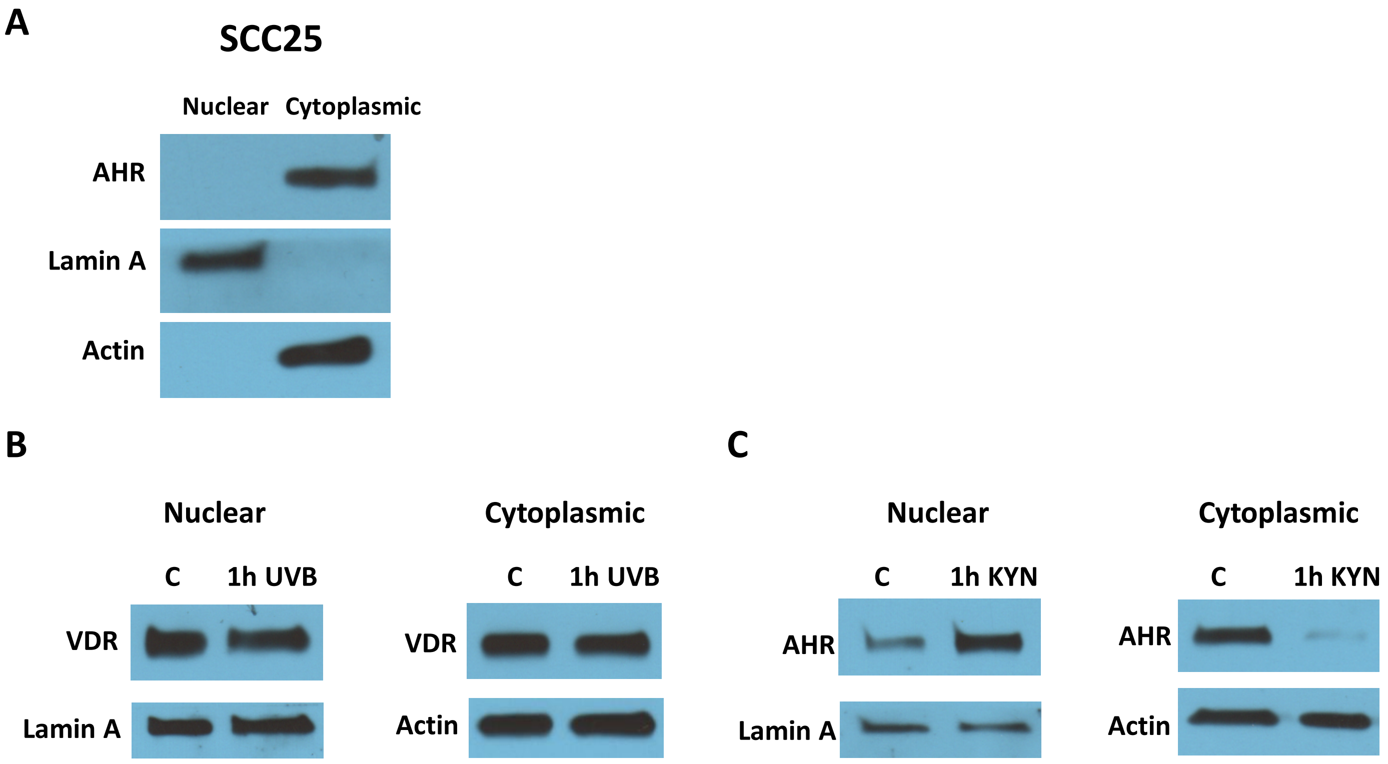


**Figure S1. A)** Western Blot analysis of distribution of nuclear and cytoplasmic proteins. Actin was probed as a cytoplasmic marker; lamin A was probed as a nuclear marker. AHR and the internal controls were taken from the same blot. Blot images are provided in the Supplementary Fig S5A. **B)** Western blot analyses of VDR protein in nuclear and cytoplasmic fractions in SCC25 cells 1 hr following irradiation with UVB (15 min). VDR and the internal controls were taken from the same blot. Blot images are provided in the Supplementary Fig S5B. **C)** Western Blot analyses of AHR from nuclear and cytoplasmic fractions in SCC25 cells, 1 hr following treatment with 50 uM Kynurenin. AHR and the internal controls were taken from the same blot. Blot images are provided in the Supplementary Fig S5C.

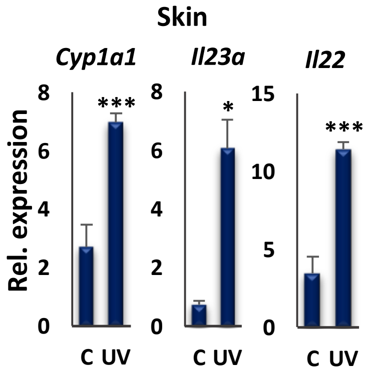


**Figure S2.** Effect of a single 15-min dose of cutaneous UVB irradiation on expression of AHR target genes *Cyp1a1, Il23a*, and *Il22* in mouse skin (n = 3 per group). RNA was extracted 4hr after UVB exposure.*P ≤ 0.05, ***P ≤ 0.001 as determined by one-way ANOVAs followed by Tukey's post hoc test for multiple comparisons.

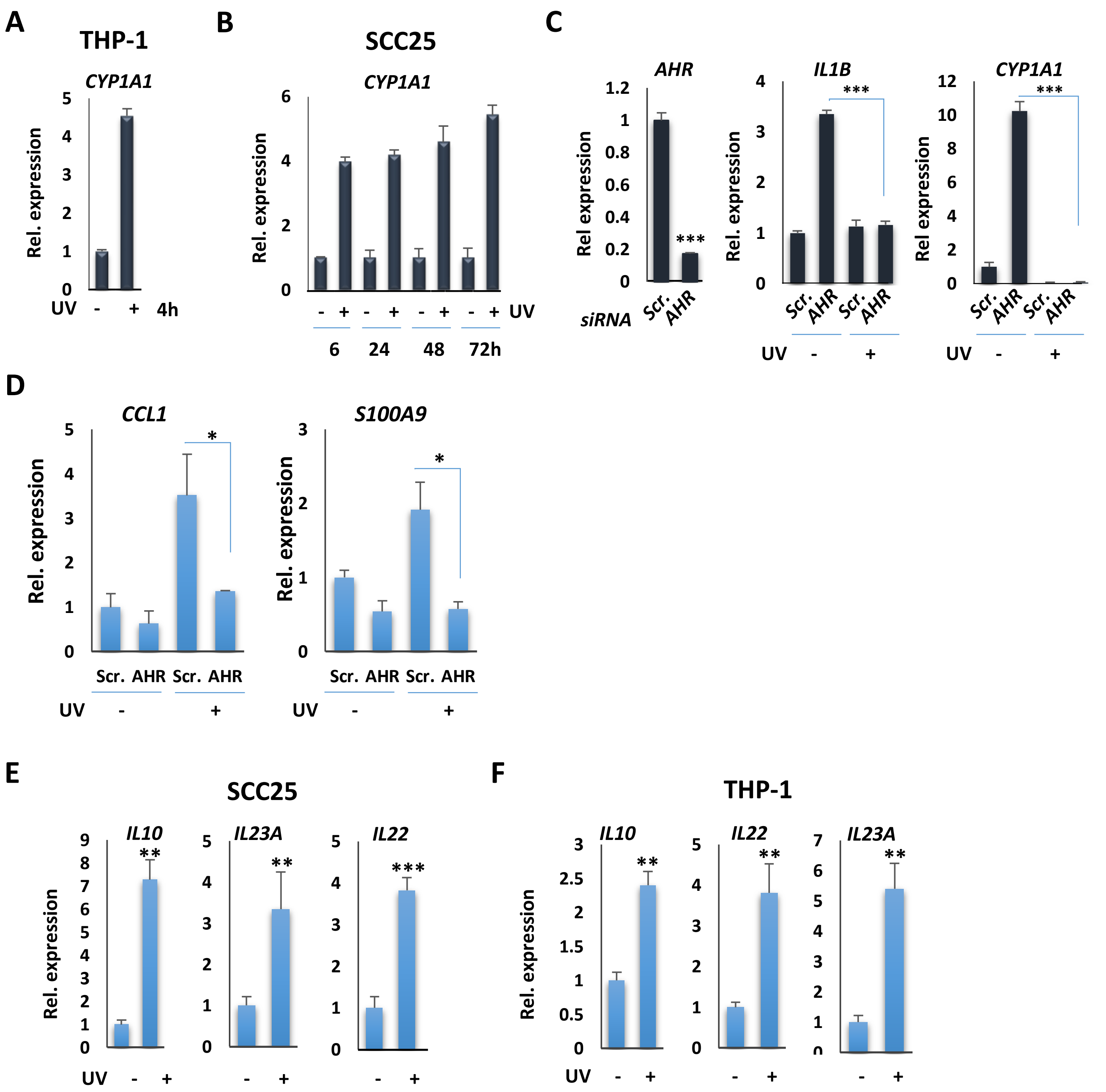


**Figure S3: A)** RT-qPCR analysis of *CYP1A1* transcription in THP-1 cells 4 hr following UVB exposure (15 min). **B)** RT-qPCR analysis of *CYP1A1* transcription in SCC25 cells 6, 24, 48, and 72 hr following irradiation with UVB (15 min). The expression of CYP1A1 in UV-exposed cells is a fold expression relative to the control cells, and is set as 1 in each time point. **C)** (left panel) RT-qPCR assay of *AHR* mRNA expression after knockdown of its gene with pooled siRNA #2 in SCC25 cells. (center and right panel) RT-qPCR analysis of *CYP1A1* and *IL1B* transcription in SCC25 cells following knockdown of the *AHR* gene and 4 hr after exposing cells to UVB for 15 min. **D)** RT-qPCR analysis of CCL1 and S100A9 transcription in SCC25 cells following knockdown of AHR expression and 4 hr after exposing to UVB for 15 min. **E)** RT-qPCR assay of *IL10*, *IL23A* and *IL22* mRNAs in SCC25 cells, 4 hr after 15 min of UVB irradiation. **F)** RT-qPCR assay of *IL10*, *IL23A* and *IL22* mRNAs in THP-1 cells, 4 hr after 15 min of UVB irradiation. $*P \leq 0.05$, $**P \leq 0.01$, $***P \leq 0.001$ as determined by one-way ANOVAs followed by Tukey's post hoc test for multiple comparisons.

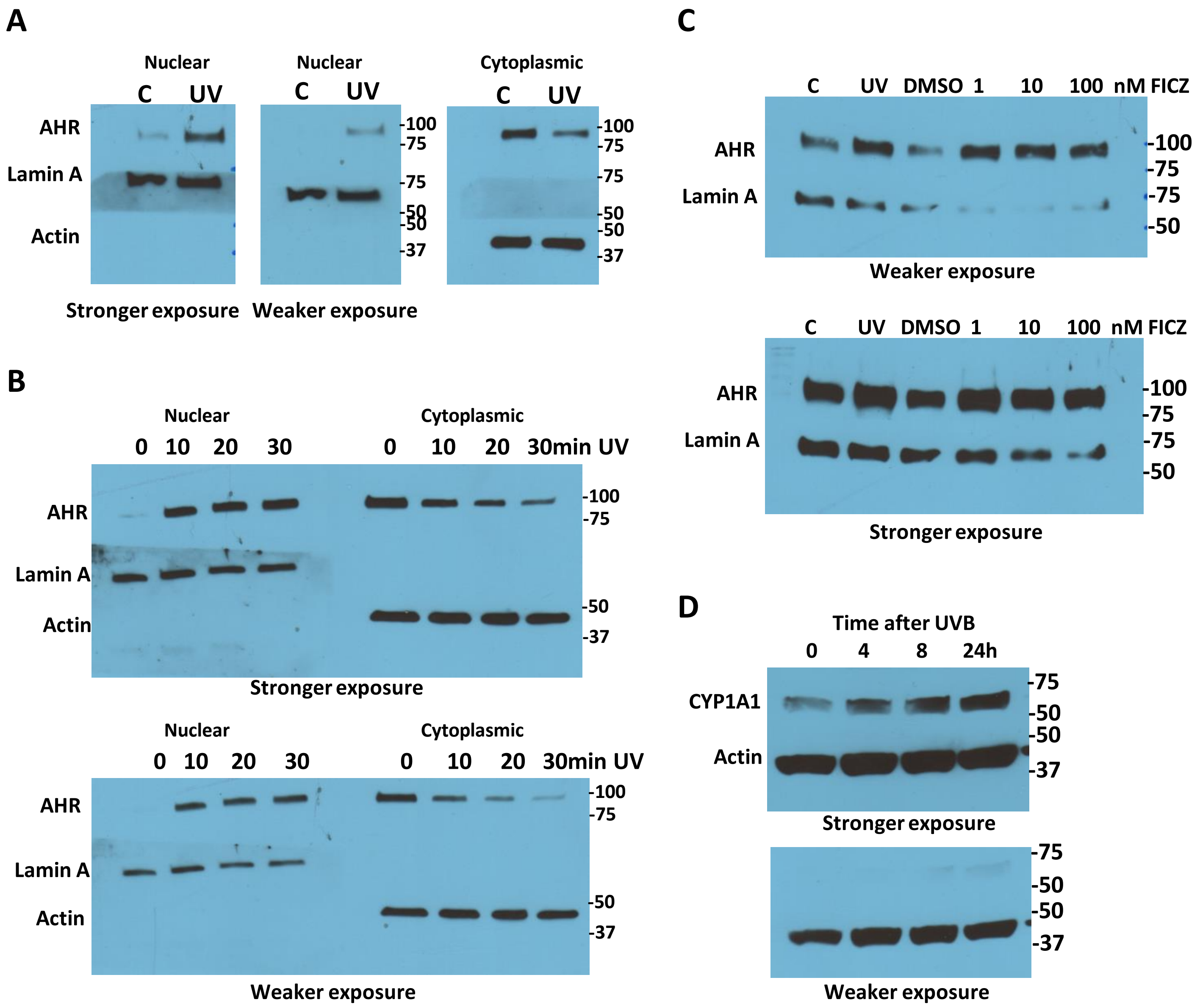


**Figure S4: A)** The blot corresponding to figure 1A. The blot was cut in three, up the 75 and 50 KD ladder bands. **B)** The blot corresponding to figure 1B. The blot was cut in two, up the 75 KD ladder band in the nuclear fraction part; and was cut in two, up the 50 KD ladder band in cytoplasmic part. **C)** The blot corresponding to figure 1C. The blot was cut in two, up the 75 KD ladder band. **D)** The blot corresponding to figure 1E. The blot was cut in two, at the 50 KD ladder band.

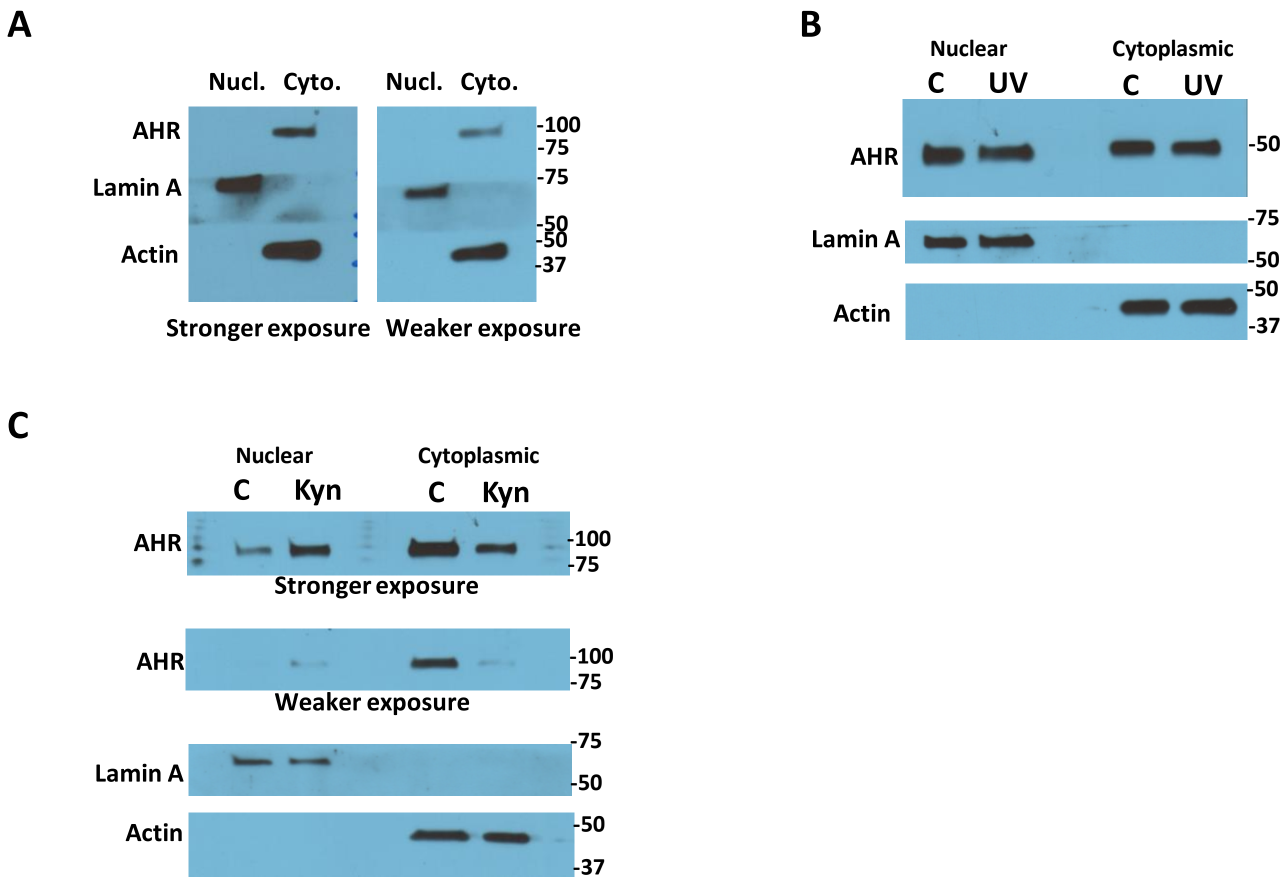


**Figure S5: A)** The blot corresponding to figure S1A. The blot was cut in three, up the 75 and 50 KD ladder bands. **B)** The blot corresponding to figure S1B. The blot was first probed for VDR and then stripped and cut in two, up the 75 and 50 KD ladder bands and probed for Lamin A and Actin. **C)** The blot corresponding to figure S1C. The blot was cut in three, up the 75 and 50 KD ladder bands.

| Table S1: Primers for gene expression analysis | | |
|---|---|---|
| Human AHR | F: CTTCCAAGCGGCATAGAGAC | R: AGTTATCCTGGCCTCCGTT |
| Human IL1B | F: AGATGAAGTGCTCCTTCCAGG | R: GGTCGGAGATTCGTAGCTGG |
| Human CYP1A1 | F: CTTGGAACCTTCCCTGATCCTTG | R: GATCTTGGAGGTGGCTGAGGTA |
| Human actin (ATCB) | F: CACCTTCACCGTTCCAGTTTT | R: AACCTAACTTGCGCAGAAAACAA |
| Human RNA18S5 | F: ACCCGTTGAACCCCATTCGTGA | R: GCCTCACTAAACCATCCAATCGG |
| Mouse Hmbs | F: GTGCCTACCATACTACCTCCTG | R: ACTCTCCTCAGAGAGCTGGTTC |
| Mouse Ahr | F: TGATGCCAAAGGGCAGCTTA | R: TGAACTGGTACCCCGATCCT |
| Mouse Il23a | F: ACCAGCGGGACATATGAATCT | R: AGACCTTGGCGGATCCTTTG |
| Mouse Il22 | F: CTGTTGACACTTGTGCGATCTCTG | R: TTGACGGGCAGCGCATTTG |
| Mouse Nqo1 | F: GATGGGAGGTACTCGAATCTGAC | R: AGCTCACCTGTGATGTCATTTCT |
| Mouse Cyp1a1 | F: TTTAAAACACGCCCGCTGTG | R: CAGGCACAATGTCCCAGGAT |
| **Primer sequences used for ChIP analysis:** | | |
| Human AHR on IL23A | F:GTCAGTTGTAGCCCTGGATGTA | R:TTGGGTAGGAAGAAGGGTTGGT |
| Human AHR on CYP1A1 (ref 1) | F: GCGCGAACCTCAGCTAGT | R:TTCCCGGGGTTACTGAGTC |
| Human AHR on IL10 (ref 2) | F: GTCTTGGGTATTCATCCCAGGTTGGGG | R:CTGTGGGTTCTCATTCGCGTGTTCCTA |
| Mouse AHR on Cyp1a1 (ref 3) | F: TATCCGGTATGGCTTCTTGC | R: CACCTTCAGGGTTAGGGTGA |
| Mouse AHR on Il22 | F: ACAGTGATTTTCATGACTTCGCGTTCT | R: TCCCAGATAGCACCTGACAACTAGACT |
| Mouse AHR on Ahr | F: GTGTGTGCGCTCCCTTTGAC | R: GAGTCCGTCCACCAGTTCGTC |


1. Vorrink, SU et al. (2013) Hypoxia perturbs aryl hydrocarbon receptor signaling and CYP1A1 expression induced by PCB 126 in human skin and liver-derived cell lines. *Toxicology and Applied Pharmacology* **274**, 408-416.
2. Gandhi R et al. (2010) Activation of the aryl hydrocarbon receptor induces human type 1 regulatory T cell–like and Foxp3+ regulatory T cells. *Nature Immunology* **11**, 846–853.
3. Amenya HZ et. al. (2016), Dioxin induces Ahr-dependent robust DNA demethylation of the Cyp1a1 promoter via Tdg in the mouse liver. *Scientific Reports* **6**, 34989.
